# Transfer of CRISPR-like activity of MIMIVIRE into Bacteria

**DOI:** 10.1101/697250

**Authors:** Azza Said, La Scola Bernard, Levasseur Anthony, Perrin Pierre, Chabrière Eric, Raoult Didier

## Abstract

MIMIVIRE is a defence system utilized by lineage A Mimiviruses against Zamilon virophages. It is composed of a helicase, a nuclease and a gene of unknown function here named *trcg* (for Target Repeat-Containing gene), which contains four 15-bp repeats identical to the Zamilon sequence. Their silencing restored susceptibility to Zamilon, and the CRISPR-Cas4-like activity of the nuclease was recently characterised. We expressed these 3 genes after transformation of a modified strain of *Escherichia coli* made resistant to ampicillin, chloramphenicol and tetracycline. The virophage repeats were replaced with four repeats of 15 nucleotides identical to a sequence in the tetracycline resistance gene. The induction of the MIMIVIRE genes restored *E. coli* sensitivity to tetracycline; the tetracycline operon and its supporting plasmid harbouring the chloramphenicol resistance gene vanished. We therefore efficiently transferred the defence system MIMIVIRE from giant Mimivirus against virophage to *E. coli* to clear it from a plasmid.

## Text

The first mechanism of defence for organisms is the cannibalization of alien sequences to prevent their multiplication^1-4^. This phenomenon has become critical in vertebrates, where the number of integrated retroviruses reaches several thousand per organism^5^, and CRISPR regulates the integration of alien gene sequences in bacteria and archaea as a defence mechanism^6^. In Mimivirus, the specific resistance of lineage A to the Zamilon virophage (a virus that infects Mimivirus) has led us to look for cannibalized sequences in an operon, which we described under the name “MIMIVIRE”^1^ (MIMIvirus VIrophage Resistance Element). Silencing 3 genes from the MIMIVIRE operon encoding a helicase gene, a nuclease gene and *trcg* (containing 4 small repeats of the virophage target) abolished MIMIVIRE activity. We proposed that this is an adaptive defence system, a proposal that has been controversial^7^. Nuclease and Mimivirus helicase have already been expressed to identify their roles^6^, and a recent new study found that the nuclease is a functional homologue of the CRISPR-Cas4 protein with dual nuclease activities^3^. Here, we explore the possibility of transferring this system into *Escherichia coli* by targeting a plasmid containing a tetracycline resistance gene.

To confirm the role of MIMIVIRE, we transformed *E. coli* with 2 plasmids, with one containing the ampicillin resistance gene and MIMIVIRE and one containing tetracycline and chloramphenicol resistance genes. We inserted the first system, including the 3 genes involved in MIMIVIRE activity, into the expression vector PP37 under the control of the IPTG-inducible T7 promoter (Figure 1). The *trcg* sequence was modified to target the tetracycline resistance gene (carried by the second plasmid), with 4 repeats of 15 nucleotides of Zamilon replaced by 4 repeats of 15 nucleotides specific to the tetracycline resistance gene (CGGCTCTTACCAGCC). The helicase and nuclease genes were added following the modified *trcg* sequence and organized into an operon under the control of the inducible T7 promoter (Figure 1). The results (Figure 2) show that transformed *E. coli* grow on tetracycline and chloramphenicol agar, but the induction of MIMIVIRE by IPTG reverses the resistance. This result was reproduced 4 times, as shown in Figure 2. We also tested the induction of the MIMIVIRE system in a liquid medium. Two colonies of *E. coli* selected on agar plates containing ampicillin and tetracycline were picked and cultured in 2 ml of medium containing ampicillin and tetracycline. Two hours later, each culture was divided into two tubes, with one supplemented with IPTG (to induce MIMIVIRE protein expression) and sampled regularly to be tested on agar containing ampicillin and tetracycline. As shown in Figure 3, the expression of MIMIVIRE causes the death of bacteria after 4 hours, with 30 and 20 times fewer colonies, confirming that MIMIVIRE abolishes tetracycline resistance. The results obtained with chloramphenicol instead of tetracycline in the above transformations show that MIMIVIRE activation also reverses chloramphenicol resistance and confirmed that the entire plasmid was destroyed, mimicking the effect of MIMIVIRE on Zamilon.

**Figure 1.**
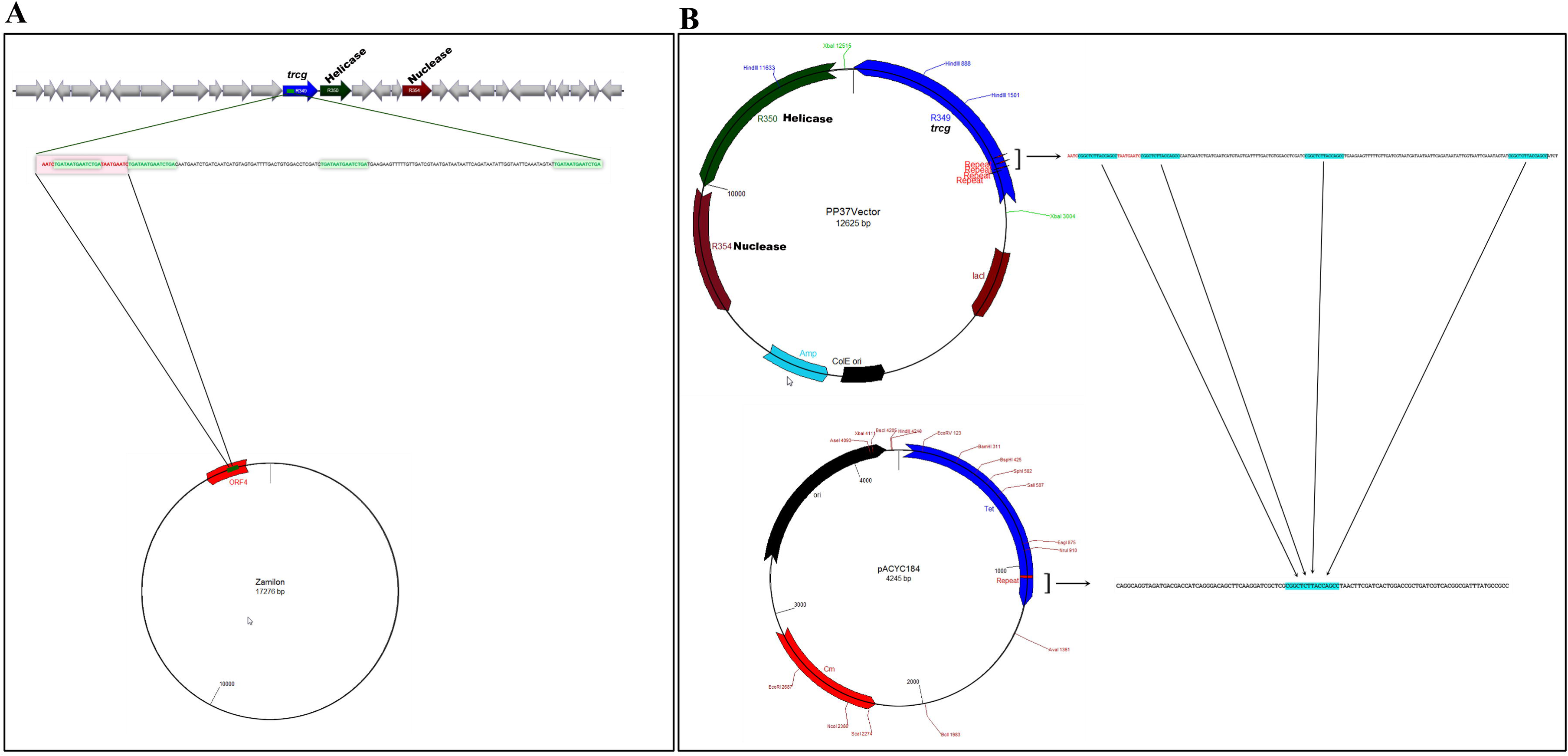
Vector construction of the prokaryotic MIMIVIRE system based on the MIMIVIRE defence system in lineage A *Acanthamoeba polyphaga* mimivirus (APMV). **A**. Nucleic acid-based immunity in MIMIVIRE against virophage Zamilon infection. The chromosomal environment of the 3 genes involved in MIMIVIRE activity (*trcg* containing 4 repeat units, helicase and nuclease genes). The 15 nucleotide repeat unit (TGATAATGAATCTGA) is specific to Zamilon ORF4. **B.** MIMIVIRE vector (PP37 vector) directed against the tetracycline resistance gene carried by the pACYC184 plasmid. The *trcg* sequence was modified to target the tetracycline resistance gene by replacing the 15 nucleotides of the Zamilon repeat with 15 nucleotides specific to the tetracycline resistance gene (CGGCTCTTACCAGCC).

**Figure 2.**
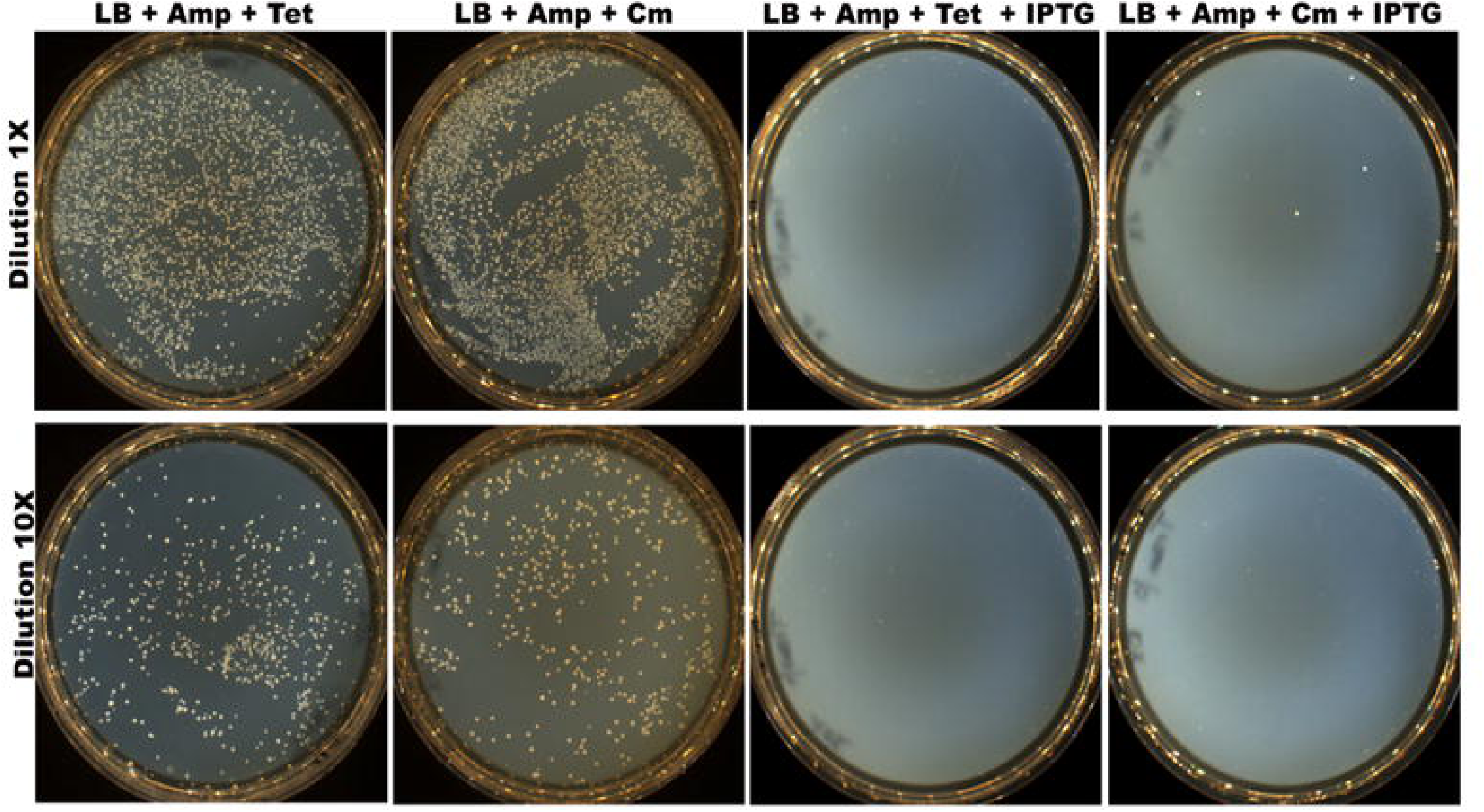
Transformation of *E. coli* harbouring the pACYC184 plasmid with the PP37 vector containing an inducible MIMIVIRE system. The bacteria are spread on LB agar + Amp + Tet or Cm with or without 1 mM IPTG to induce MIMIVIRE protein expression. Each plate was digitalized on a Scan® 1200 (Interscience, France).

**Figure 3.**
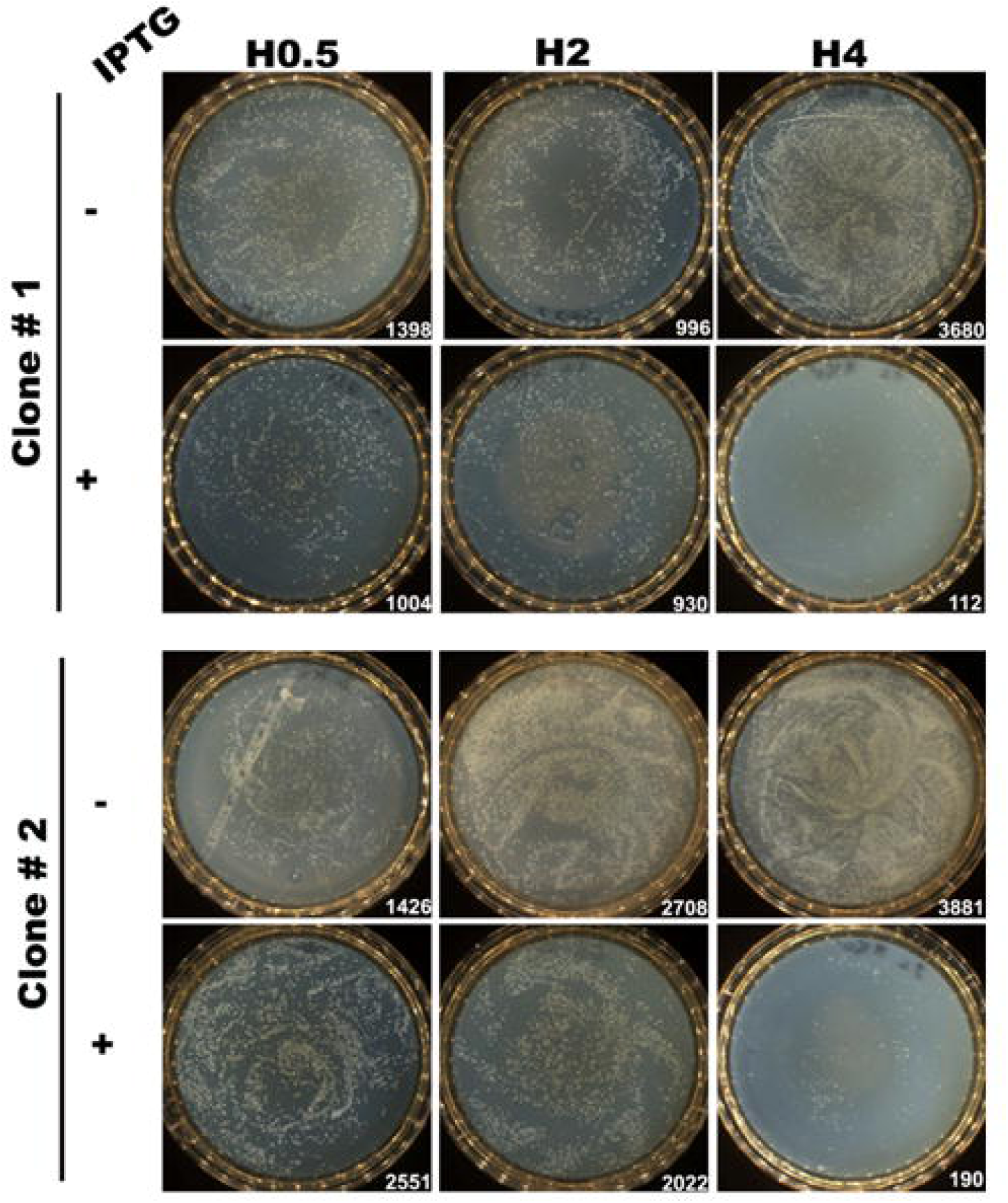
Results of the induction of the MIMIVIRE system in liquid medium. Two colonies of *E. coli* harbouring both plasmids (PP37 vector and pACYC184) were picked and cultured in 2 ml of LB + Amp + Tet medium at 37 °C under shaking conditions at 200 rpm. Two hours later, each culture was divided into two tubes, in which one was supplemented with 1 mM IPTG to induce MIMIVIRE protein expression. After 30 minutes (H0.5), 2 hours (H2) and 4 hours (H4), 10 µl of cell culture was diluted into 1 ml, from which 100 µl was spread on LB + Amp + Tet agar plates. The next day, each plate was digitalized on a Scan® 1200 (Interscience, France), and colonies were counted. The number of colonies is shown below each plate.

To exclude possible lethal MIMIVIRE expression, we tested the viability of induced bacteria on selective media. Four colonies of *E. coli* growing on ampicillin were tested under the same conditions, and the *E. coli* culture was not hampered, confirming that the expression of MIMIVIRE was not toxic by itself (Figure 4). Our results prove that MIMIVIRE can be transferred into *E. coli* and act against a new gene by including 15 specific nucleotide repeats of the targeted sequence.

**Figure 4.**
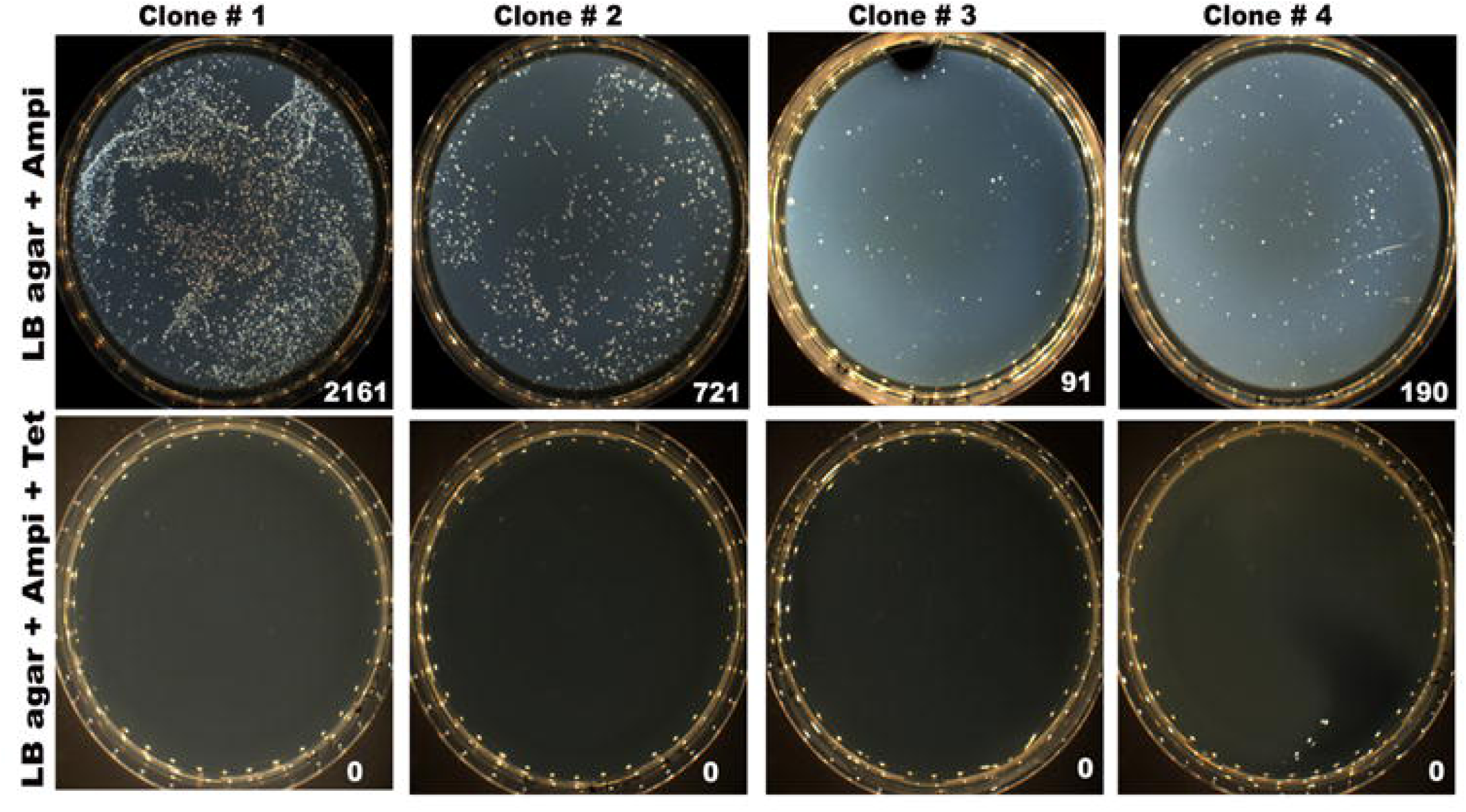
Lack of lethal effect of MIMIVIRE expression. Four colonies of *E. coli* harbouring both plasmids (PP37 vector and pACYC184) were picked and cultured in 1 ml of LB + Amp + Tet medium at 37 °C under shaking conditions at 200 rpm. Two hours later, each culture was supplemented with 1 mM IPTG to induce MIMIVIRE protein expression. After 4 hours, 10 µl of cell culture was diluted into 1 ml, from which 100 µl was spread on LB + Amp + Tet agar plates or LB + Amp agar plates. The next day, each plate was digitalized on a Scan® 1200 (Interscience, France), and colonies were counted. The number of colonies is shown below each plate.

In conclusion, we show herein that the MIMIVIRE system of resistance to virophages may be exported into bacteria and acts as CRISPR-Cas molecular scissors despite its different organization. We believe that the use of 4 repeats mechanically increased the probability of generating heteroduplex DNA-RNA, which inhibits DNA polymerase progression. We hypothesize that the helicase opens the heteroduplex DNA-RNA and then the nuclease digests single-strand DNA^3^. In any case, we proved the activity of a new defence mechanism expressed in bacteria that may add to our arsenal in modifying eukaryotic and cell genomes.

## Methods

### 1. Plasmid design and construction

The 3 genes involved in MIMIVIRE activity were codon-optimized for *E. coli* expression, synthesized by GenScript and cloned into pET24b(+) under the control of the IPTG-inducible T7 promoter (Figure 1). The *trcg* sequence was modified to target tetracycline resistance by replacing the 15 nucleotides of the Zamilon repeat with 15 nucleotides specific to the tetracycline resistance gene (CGGCTCTTACCAGCC). To identify this sequence, the tetracycline resistance gene was fragmented with a sliding window of 15 nucleotides and a step of one nucleotide to generate all possible 15 nucleotide long sequences, which were used as queries to search BLASTn for similar sequences in the two vector sequences (PP37 vector, 12,565 bp; pACYCl84 with the tetracycline resistance gene sequence deleted, 4,245 bp) and then submitted to a BLAST search against the *E. coli* strain BL21 (DE3) genome sequence downloaded from the NCBI GenBank database 55 (NC_ 012971.2). Two fragments were identified that had the lowest similarity with these sequences. One of them, CGGCTCTTACCAGCC, was used to construct the PP37 vector. The helicase and nuclease genes were added following the modified *trcg* and organized into an operon under the control of the inducible T7 promoter (Figure 1).

### 2. Transformation assay

The plasmids used in these experiments were PP37 vector synthesized by GenScript, allowing inducible expression of the MIMIVIRE system under T7 promoter control (Figure 1) and pACYC184 (MoBiTec company, Ref: V32402). *E. coli* One Shot BL21 (DE3) chemically competent cells (Thermo Fisher, Ref: C600003) were transformed with the pACYC184 plasmid according to standard protocol and the manufacturer’s instructions. Bacteria were spread on LB agar with the appropriate antibiotic (12 µg/ml of tetracycline and 100 µg/ml of ampicillin). Bacteria were spread on a tetracycline (Tet) selective LB agar plate (day 1). The day after, one clone was picked and cultured in 10 ml of Tet selective LB medium overnight at 37 °C under shaking conditions at 200 rpm. On day 3, 200 µl of cell culture was used to inoculate 20 ml of fresh Tet LB medium until the OD at 600 nm reached 0.7. Bacteria were harvested by centrifugation at 7000 rpm for 1 minute at 4 °C and washed three times with 10% glycerol. After the first centrifugation step, all steps were performed on ice in a cold room. Bacteria were resuspended in 50 µl of 10% glycerol, and 50 ng of PP37 vector was added. The mix was transferred to a Gene Pulser cuvette with a 0.1 cm gap (Bio-Rad; Ref 165-2089), and an electro-pulse was delivered with a MicroPulser Electroporator (Bio-Rad; Ref: 165-2100) using the Ec1 program. The electroporation time indicated by the device after the pulse was 5.7 ms. After electroporation, 1 ml of LB was added immediately, and the cells were incubated for 1 hour at 37 °C with shaking at 200 rpm. *E. coli* BL21 (DE3) harbouring both plasmids (PP37 vector and pACYC184) were selected in LB agar supplemented with 12 µg/ml of tetracycline and 100 µg/ml of ampicillin.

### 3. Induction of the MIMIVIRE system in *E. coli*

To test the effectiveness of the MIMIVIRE system in *E. coli* harbouring both plasmids (PP37 vector and pACYC184), several colonies selected from LB agar plates containing ampicillin and tetracycline (100 μg/ml and 12 μg/ml, respectively) were picked and cultured in 2 ml of LB medium containing ampicillin and tetracycline (100 μg/ml and 12 μg/ml, respectively) at 37 °C with shaking at 200 rpm. Two hours later, each culture was divided into two tubes, with one supplemented with 1 mM IPTG to induce MIMIVIRE protein expression. At regular time intervals, 10 µl of cell culture was diluted to 1 ml, from which 100 µl was spread on LB agar plates containing ampicillin and tetracycline (100 μg/ml and 12 μg/ml, respectively). The next day, each plate was digitalized on a Scan® 1200 instrument (Interscience, France), and colonies were counted according to the manufacturer’s recommendations. As a negative control to verify that bacterial death was not due to induction only, the same experiment was performed on agar plates without tetracycline.

## Acknowledgements

This work was supported by the French Government under the “*Investissements d’avenir*” programme managed by the *Agence Nationale de la Recherche (ANR)*, [reference: *Méditerranée-Infection* 10-IAHU-03], by Région Provence-Alpes-Côte d’Azur and European funding FEDER PRIMI.

## Author contributions

SA provided technical manipulation and redaction

BL provided concept and redaction

AL provided performed metagenomic analysis

PP performed the manipulations

EC provided concept

DR conceived the study and designed the methodology and wrote the manuscript

## References

1. B. La Scola, et al., “The virophage as a unique parasite of the giant mimivirus,” Nature 455(7209), 100 (2008).

2. A. Levasseur, et al., “MIMIVIRE is a defence system in mimivirus that confers resistance to virophage,” 531(7593), 249 (2016).

3. C. Dou, et al., “Structural and Mechanistic Analyses Reveal a Unique Cas4-like Protein in the Mimivirus Virophage Resistance Element System,” Cell 3, 1 (2018).

4. P. Colson, et al., “HIV infection en route to endogenization: two cases,” Clin. Microbiol Infect. 20(12), 1280 (2014).

5. K. S. Makarova, et al., “An updated evolutionary classification of CRISPR-Cas systems,” Nat. Rev Microbiol 13(11), 722 (2015).

6. M. Bekliz, et al., “[The defence system MIMIVIRE in mimivirus illustrates Red Queen hypothesis],” Med. Sci (Paris) 32(10), 818 (2016).

7. J. M. Claverie and C. Abergel, “CRISPR-Cas-like system in giant viruses: why MIMIVIRE is not likely to be an adaptive immune system,” Virol. Sin. 31(3), 193 (2016).

